# Estimating resource acquisition and water-use traits in wine grapes using reflectance spectroscopy

**DOI:** 10.1101/2025.10.29.685438

**Authors:** Rachel O. Mariani, Marney E. Isaac, Kimberley Cathline, Gavin Robertson, Adam R. Martin

**Affiliations:** Department of Physical and Environmental Sciences, University of Toronto Scarborough, Canada; Horticultural & Environmental Sciences Innovation Centre, Niagara College, Canada; Food and Beverage Innovation Centre, Niagara College, Canada

**Keywords:** agroecology, functional traits, plant hydraulic traits, high-throughput phenotyping, reflectance spectroscopy

## Abstract

In agroecosystems, the variable expression of crop functional traits is expected to play a role in key processes, including plant nutrient cycling and water acquisition, that confer ecosystem resistance and/ or resilience to environmental change. The ability to estimate crop trait data is therefore critical to predict crop responses to environmental change, enabling more informed diagnosis of crop performance and on-farm management strategies. Yet, many traditional methods for quantifying plant traits are time-consuming and resource-intensive, limiting sample sizes and study durations. In response, high-throughput phenotyping— specifically reflectance spectroscopy— has emerged as a key element of plant trait research, capable of estimating plant traits more rapidly. However, little is known about whether or not reflectance spectroscopy can detect within-species variation in resource acquisition and plant-water traits. Using wine grapes (*V. vinifera* subsp. *vinifera*) as a focal crop, this study aimed to assess the ability of reflectance spectroscopy and the subsequent partial least squares regression modelling approach to quantify intraspecific variation in 12 functional traits across 12 different cultivars. Results showed significant differences in traits, especially in the photosynthetic and hydraulic traits, among closely related cultivars, falling along a resource-conservative to resource-acquisitive axis of variation. We also found that reflectance differentiated this fine-scale trait variation, specifically in leaf chemical and morphological traits, contributing to higher accuracy, and indicating that this HTP approach is viable for detailed trait estimation in diverse agroecosystems.

## Introduction

In agricultural systems, environmental change driven by deviations in temperature and precipitation patterns is exerting profound impacts on crop growth and productivity (1,2). While observations and predictions vary widely across crop species and geographies, recent studies have reported anywhere between 7-23% declines in crop yield (1). Such changes largely represent the impact of fluctuations in precipitation and changes in growing and non-growing season temperatures on crop phenology, ecophysiology, and nutrient uptake (3). These complex interactions between environmental impacts and plants scale up to influence crop performance, yield, and quality on farms (e.g., 2–4).

Globally, wine grapes are one of the world’s most valuable crops, and the regions of grape production are one of the major drivers of crop quality and value (2). In the province of Ontario, Canada, for example—the region of our study—wine grapes and wine are among the highest value-added agricultural products, with a $5.49 billion total economic impact in 2024-25 (5). However, the suitability of the world’s wine-growing regions is expected to shift in the coming years and decades. Rising temperatures and prolonged droughts may hinder production, making hot and dry regions less suitable for cultivation. Meanwhile, mid-latitude wine-growing regions could experience a higher risk of spring frosts and increases in precipitation, which are expected to impact plant phenology with negative consequences for production (2). Wine grapes are vulnerable to environmental change, as factors such as temperature variation, precipitation patterns, and extreme weather events can influence grapevine growth and fruit quality (4,6).

Projected shifts in precipitation patterns due to climate change pose significant challenges in viticulture. Predictions suggest that under a 2 °C global temperature increase scenario, up to 56% of all current wine-growing regions will become unsuitable for growing due to changes in grapevine phenology and grape composition, with this number increasing to 85% with a 4 °C increase (2,7). Thus, wine growers will need to consider adaptation strategies to ensure consistency in production and quantity under variable environmental conditions (2,8). More specifically, varietal selection and cultivar turnover are potentially key climate change mitigation strategies to apply to ensure vine resilience to changes in growing regions. The turnover relies on varietal selection, where grapevine varieties are chosen based on their adaptability and used in various combinations of scion, rootstock, and training systems (9). Wine grapes show significant genomic and phenotypic diversity due to years of artificial selection, leading to cultivars with traits that aid in environmental adaptation. Trait variation in leaves related to photosynthesis, gas exchange, and thermoregulation exhibits wide morphological, chemical, and physiological differences that are influenced by both environmental and genetic factors (9,10).

Plant responses to environmental change drivers can be multifaceted and governed by synchronized and/ or independent leaf, root, whole-plant, and other stress response traits (Rezaei et al., 2023). One key hypothesis in ecosystem science is that inter- and intra-specific variation in leaf functional traits—the characteristics of plants that determine responses to the surrounding environment —play a role in governing plant- and ultimately ecosystem-level responses to climate change (11–13). Generally, the literature on functional traits underpins an expectation that greater functional trait variation leads to greater ecosystem resistance and/ or resilience to environmental change (14–16). While much of the earlier work in this area focused on interspecific trait variation, more recently, research suggests that higher intraspecific trait variation can also indicate stronger resistance and adaptation of plant communities to change (14).

Quantifying intraspecific trait variation is particularly important in agroecosystems, where a limited number of plant species, varieties, or cultivars dominate and ultimately influence ecosystem functioning due to their high abundance; however, many “traditional” methods for this are often time-consuming and resource-intensive, which in turn limits researchers to smaller sample sizes and/ or shorter-term studies (17,18). In response to challenges associated with plant trait data collection, high-throughput phenotyping (HTP) has emerged as vital in plant trait research. Compared to “traditional” data collection methods, HTP tools can rapidly quantify several plant traits in a field setting (18).

At the ecosystem level, remote and hyperspectral sensing are key components of HTP approaches. On the plant- or leaf-level, reflectance spectroscopy is a technique being developed and employed to assess interspecific plant trait variation (19,20), especially for plant traits, such as specific leaf area (SLA), photosynthetic capacity, and leaf carbon: nitrogen ratios. Earlier research primarily focused on quantifying interspecific trait variation among plants of different functional types, especially those with diverse evolutionary histories (21). Reflectance spectroscopy has proven effective in explaining trait variation, especially for leaf morphological and certain physiological traits, thus indicating that this technique is suitable for rapid trait assessment in natural and managed ecosystems. However, recent interest in intraspecific trait variation highlights the need to better understand and quantify the ability of reflectance spectroscopy to quantify trait variation within species (16,20,21).

Numerous studies have investigated plant water status using leaf optical properties in recent decades, and the majority have focused on leaf structural or chemical traits—only a few studies have explored the potential of reflectance spectroscopy to estimate within-species variation in leaf water potential (21,22). To the best of our knowledge, there are limited studies that evaluate isotopic signatures (δ ^13^C) and predawn water potential (Ψ_pd_), traits that are critical indicators of plant water relations and water-use strategies (23,24) These traits help assess drought tolerance by providing measures of stomatal behaviour and plant water status, which in turn influence key physiological processes such as plant stomatal conductance (*g*_s_) and maximum photosynthetic activity (*A*_max_) under periods of environmental stress. The use of spectroscopy to estimate δ ^13^C and Ψ_pd_ will make assessing the resilience of wine grape cultivars more feasible (23–25).

This study aims to determine intraspecific leaf traits associated with plant-water relations and leaf resource acquisition across multiple wine grape cultivars to assess the following: 1) do wine grape cultivars differ in their leaf traits, specifically in their leaf morphological, chemical, physiological, and water-use traits; and if so, 2) can reflectance spectroscopy quantify this intraspecific trait variation. We hypothesize that each wine grape cultivar will exhibit significant variation in leaf traits, reflecting their genetic variation, especially in their physiological and water-use traits. Additionally, we hypothesize that reflectance spectroscopy can reliably quantify intraspecific variation in leaf traits among wine grape varieties, with spectral signatures capturing key properties associated with morphological, chemical, physiological, and water-use traits, as seen in previous studies using different wine grape varieties (26).

## Materials and Methods

### 2.1 Study design and trait collection

This study was conducted at the Niagara College Teaching Vineyard in Niagara-on-the-Lake, Ontario (43.146237 °N, -79.156810 °W). The study site is an operational teaching vineyard containing multiple cultivars of wine grapes (*V. vinifera* subsp. *vinifera*) distributed across the 16.2-ha farm in rows separated by either cover crops or unmanaged grasses. The soil at the vineyard is a mineral-rich mix of sand, gravel, loam, and red clay, which creates a substrate ideal for wine grape production. At the site, 12 different wine grape cultivars belonging to seven different varieties of both red and white grapes are present, including Riesling clones 23 and 171 (R23 and R171), Pinot Noir clones 89 and 828 (PN89 and PN828), Merlot clones 384 and 181 (M348 and M181), Cabernet Sauvignon clones 29 and 412 (CS29 and CS412), Cabernet Franc clones 327 and 314 (CF327 and CF314), Sauvignon blanc clone 906 (SB906) and Viognier clone 642 (V642). All cultivars are grown on a Selection Oppenheim (SO4) rootstock except Viognier 642, which is grown on the Millardet et de Grasset (101–14) rootstock. The vine training system is primarily a vertical shoot positioning (VSP) system, which is most commonly used for varieties that thrive under improved sunlight exposure. The vineyard is not irrigated, though it is tile-drained, allowing for better soil aeration.

For each clone, five vines in three different planting rows equally spaced along the northwest side of the vineyard were selected for functional trait characterization. Specifically, a total of *n*=171 leaves were sampled from *n*=15 vines per clone, equally across *n*=12 clones at the vineyard. On each vine, we selected one fully expanded, west-facing upper canopy leaf for reflectance spectroscopy and trait collection, such that all selected leaves were approximately the same size and free from any visible signs of damage.

For each sample leaf, two main sets of in-field measurements were collected. First, physiological gas exchange data were collected using an LI-6800 portable photosynthesis system affixed with a 6800-03 large light source (LICOR Biosciences, Lincoln, Nebraska, USA).

Specifically, for each leaf in our dataset the LI-6800 was used to measure stomatal conductance (*g*_s_; mmol^-2^ s^-1^), light-saturated photosynthetic capacity (*A*_max_; *μ*mol CO_2_ m^-2^ s^-1^), and evapotranspiration (*E*; mmol m^-2^ s^-1^), under the following leaf chamber conditions: CO_2_ concentrations of 420 ppm, saturating light levels of 1,500 *μ*mol of photosynthetically active radiation (PAR) m^-2^ s^-1^ (with irradiance peaks in blue [453 nm], green [523 nm], and red [660 nm] wavebands), relative humidity (RH) of 50%, leaf vapour pressure deficits (VPD) of 1.7 kPa, and leaf temperatures of 25 °C (Macklin, et al., 2022). All physiological trait measurements were taken between 7:00-13:00 to ensure the leaves were not approaching stomatal closure. To ensure the leaves acclimatized to the leaf chamber conditions, we waited approximately five minutes following application of the sensor head to the leaf before taking measurements, and only captured measurements once the leaves met predetermined stability criteria. Additionally, to ensure stomata were not closed during measurements, no measurements were taken if *g*_s_ fell below 0.08 mmol^-2^ s^-1^. Physiological traits measured in the field were used to determine intrinsic water-use efficiency (WUE_intr_; *μ*mol CO_2_ mol^-^¹ H_2_O) calculated as *A*_max_/ *g*_s_, and instantaneous water-use efficiency (WUE_inst_; *μ*mol CO_2_ mol^-^¹ H_2_O) calculated as *A*_max_/ *E*.

Following gas exchange measurements, we collected spectral reflectance data for each leaf using an SVC HR-1024i handheld spectroradiometer outfitted with an LC-RP Pro leaf clip that included a calibrated internal light source (Spectra Vista Corporation, Poughkeepsie, New York, USA). The SCV HR1024i is a full-range spectroradiometer with a range of 350–2500 nm (Spectra Vista Corporation, Poughkeepsie, New York, USA) with a spectral resolution of ≤3.5 nm (350-1000 nm), ≤9.5 nm (1000-1800 nm), and ≤ 6.5 nm (1800-2500 nm), outfitted with an LC-RP Pro leaf clip that includes a calibrated internal light source. Reflectance spectra were collected on the same adaxial side of the leaves as the gas exchange measurements with an integration time of 2 s. The reference spectra were taken on a white Spectralon standard before each measurement. These measurements were taken the day following leaf physiological measurements, in conjunction with the predawn water potential measurements.

To determine pre-dawn water potential, all leaves sampled for leaf physiological measurements and reflectance spectroscopy scans were removed from their vines before sunrise (between 03:30-05:30). Leaves were stored in darkness while maintaining leaf hydration by wrapping each sample in a saturated paper towel, sealing them in a polyethylene bag, and storing them in a cool insulated bag that blocked all sunlight until they were successfully transported.

Predawn water potential (Ψ_pd_) measurements were executed using a Portable Plant Water Potential Console (Soil Moisture Equipment Corp., Goleta, CA., USA). For each leaf, a clean blade was used to slice through the petiole across the horizontal axis, and the leaf was inserted into the chamber with a microscope camera aimed at the cut petiole to detect when sap was first exuded, at which point Ψ_pd_ was recorded.

Following measurements of Ψ_pd_, all leaves were transported to the University of Toronto Scarborough, Canada, for measurement of morphological and chemical traits. First, all leaves were weighed for leaf fresh mass (g) and scanned for fresh leaf area (cm^2^) using an LI-3100C leaf area meter (LICOR Biosciences, Lincoln, NE, USA). Subsequently, all leaves were dried at 60 °C and reweighed for dry mass (g), which was then used to calculate leaf dry matter content (LDMC) as mg of dry mass/ g of fresh mass. Following these steps, leaves were ground into fine tissue using a MM40 Retsch ball mill (Retsch Ltd., Hann, Germany), and leaf N and C concentrations were determined on ∼0.15 g of leaf tissue using a LECO CN 628 elemental analyzer (LECO Corporation, 2024). Then LMA was calculated as leaf dry mass (g)/ leaf area (m^2^). Finally, we assessed δ ^13^C on ∼1 mg of the same leaf tissue, using a Picarro G2131-i isotope and gas concentration analyser (Picarro Inc., Santa Clara, CA, USA).

### 2.2 Statistical analyses

We used R statistical software v. 4.4.3 (R Foundations for Statistical Computing, Vienna, Austria) for all data analysis. First, we calculated the summary statistics for each leaf trait and assessed data normality using both the Shapiro–Wilk test and histograms associated with base R and the “stats” package. Then, variance component estimation was completed using the “nlme” R package (27) and a nested model approach, using the following components: origin, red/white cultivars, clone, and row. Outliers were detected and removed before continuing the analysis, following the interquartile range (IQR) method using a factor of 2, as this approach is used for normally distributed data with moderate to low variability.

We then applied the methodology outlined by (28) and (26) to assess whether or not, and to what extent, reflectance spectroscopy predicts traits using a partial least squares regression (PLSR) approach. All PLSR models incorporated reflectance data from the 400–2400 nm wavelength range and were designed to predict raw and/ or log-transformed trait values. For each PLSR model, the dataset was divided into a calibration set (80% of the data) and a validation set (the remaining 20%) using the “pls” and “spectrolab” packages (29,30). Since we aimed to evaluate A) whether or not reflectance spectra can predict variation in leaf traits across grape varieties and B) how effectively the PLSR modelling approach could distinguish between cultivars, we conducted the analysis using two types of data splits. First, we split the dataset by cultivar identity, ensuring that the calibration and validation sets contained approximately equal representation of trait and spectral data. Second, we performed a fully randomized split, allowing the proportion of data from each cultivar to randomly vary.

Model performance was evaluated using the validation datasets as an external test, where predicted values were compared to the observed values in the validation data subset. For each model, we used the coefficient of determination (*r*^2^), root mean squared error of prediction (RMSE), and percent RMSE (%RMSE) as the key model evaluation criteria. Then, to further assess model performance, we analysed the model regression coefficients and variable importance in projection (VIP) scores to determine which spectral regions significantly contributed to trait prediction. Additionally, we conducted a jackknife permutation analysis using the jackknife functions in the “pls” package (30) to assess model uncertainty. The jackknife coefficients were compared to those from the full model. Finally, using both the full model and jackknife outputs, we calculated the mean predicted values along with 95% confidence and prediction intervals for each trait in the validation dataset.

## Results

### 3.1 Trait variation in wine grapes

All physiological and morphological leaf traits, and their corresponding descriptive statistics, including mean, standard deviation, range, and coefficients of variation (CV), are presented in Table 1, alongside variance components attributed to origin, red/white grape cultivars, clone, and row. Across the measured traits, physiological traits such as *E* and WUE_inst_ expressed the highest variability, with coefficients of variation of 34.8 and 31.0%, respectively, indicating potentially greater sensitivity to environmental and individual-level differences. In contrast, the leaf chemical traits, namely leaf C concentrations, showed lower variability, suggesting a more conservative response to individual-level change. Generally, physiological traits varied more widely than the morphological and chemical traits (Table 1).

**Table 1.** Descriptive statistics and variance partitioning for 12 leaf traits measured across 12 wine grape clones (n=171).

| Trait | Descriptive Statistics |  |  |  |  |  | Variance Component Estimation |  |  |  |  |
| --- | --- | --- | --- | --- | --- | --- | --- | --- | --- | --- | --- |
|  | SW | Median/ Mean | SD | Min | Max | CV | Origin | R/W | Clone | Row | Unex |
| $E$ | 0.992 | 0.0032 | 0.0013 | 0.00069 | 0.01 | 34.83 | 0.42 | <0.001 | 0.38 | 0.08 | 0.13 |
| $A_{\max}$ | 0.989 | 21.83 | 3.35 | 12.95 | 30.35 | 15.35 | 0.27 | <0.001 | <0.001 | 0.19 | 0.53 |
| $g_s$ | 0.983* | 0.26 | 0.08 | 0.10 | 0.47 | 30.98 | <0.001 | <0.001 | 0.54 | 0.15 | 0.31 |
| $WUE_{\text{inst}}$ | 0.728** | 6.70 | 2.28 | 3.59 | 28.98 | 33.98 | 0.59 | <0.001 | 0.29 | 0.04 | 0.08 |
| $WUE_{\text{intr}}$ | 0.948** | 0.09 | 0.03 | 0.04 | 0.17 | 29.32 | <0.001 | <0.001 | 0.52 | 0.15 | 0.33 |
| N | 0.993 | 2.75 | 0.35 | 1.59 | 3.62 | 12.66 | <0.001 | 0.12 | <0.001 | 0.05 | 0.83 |
| C | 0.984* | 45.81 | 0.78 | 42.57 | 47.54 | 1.70 | 0.08 | <0.001 | 0.32 | <0.001 | 0.61 |
| $\delta^{13}\text{C}$ | 0.983* | -27.88 | 1.22 | -30.86 | -22.69 | -4.37 | <0.001 | <0.001 | 0.08 | 0.08 | 0.84 |
| $\Psi_{\text{pd}}$ | 0.978* | 5.88 | 1.47 | 1.25 | 10.25 | 24.96 | 0.00 | 0.18 | 0.21 | 0.07 | 0.55 |
| Area | 0.977* | 150.74 | 34.30 | 75.31 | 289.37 | 22.76 | 0.15 | <0.001 | 0.09 | <0.001 | 0.77 |
| LDMC | 0.667** | 255.69 | 21.98 | 163.95 | 568.35 | 8.60 | <0.001 | 0.66 | 0.02 | <0.001 | 0.91 |
| LMA | 0.87** | 71.67 | 9.82 | 38.53 | 122.61 | 13.71 | <0.001 | <0.001 | 0.04 | <0.001 | 0.96 |

$E$ : evapotranspiration ( $\text{mmol H}_2\text{O m}^{-2} \text{s}^{-1}$ ); $A_{\max}$ : light-saturated photosynthetic capacity ( $\mu\text{mol CO}_2 \text{m}^{-2} \text{s}^{-1}$ ); $g_s$ : stomatal conductance ( $\text{mmol m}^{-2} \text{s}^{-1}$ ); $WUE_{\text{inst}}$ : instantaneous water-use efficiency ( $\mu\text{mol CO}_2 \text{mol}^{-1} \text{H}_2\text{O s}^{-1}$ ); $WUE_{\text{intr}}$ : intrinsic water-use efficiency ( $\mu\text{mol CO}_2 \text{mol}^{-1} \text{H}_2\text{O}$ ); N: leaf nitrogen concentration (% dry mass); C: leaf carbon concentration (% dry mass); $\delta^{13}\text{C}$ : isotopic signature (‰); $\Psi_{\text{pd}}$ : predawn water potential (kPa); Area ( $\text{cm}^2$ ); LDMC: leaf dry matter content ( $\text{mg g}^{-1}$ ); LMA: leaf mass per area ( $\text{g m}^{-2}$ ). Also shown here are descriptive statistics associated with a Shapiro–Wilk test (SW) where \*: $p < 0.05$ ; \*\*: $p < 0.001$ ; SD: standard deviation; R/W: red or white; Unex: unexplained.

For two plant traits, *E* and WUE_inst_, a significant portion of the variance in trait data was explained by cultivar origin (either from a warm or cool climate region). Clone identity explained much of the variation in *g_s_* (0.54), Ψ_pd_ (0.21). WUE_intr_ (0.52) leaf C (0.32), and WUE_inst_ (0.29) (Table 1). Clone identity was the second most important source of trait variation; however, a majority of the traits expressed a significant amount of ‘unexplained variance’, meaning that there was a large portion of the variability of these traits that was not captured in our analysis or by any of our explanatory factors analysed here.

### 3.2 Trait variation across cultivars

All cultivars exhibited similar patterns in both physiological and morphological traits (Table 2, Figure 1). Across cultivars, photosynthetic traits, namely *E*, *A_max_*, and *g_s_*, expressed significant variation. Cultivars such as SB906, R23, and PN828 exhibited high maximum assimilation rates (22.3±0.79, 21.1±0.51, and 21.5±0.67 *μ*mol CO_2_ m^−2^ s^−1^, respectively) and stomatal conductance values (0.32±0.01, 0.28±0.02, and 0.26±0.01 mmol^−2^ s^−1^, respectively); however, M181 and M348 presented lower values, reflecting lower photosynthetic capacity.

**Figure 1.**
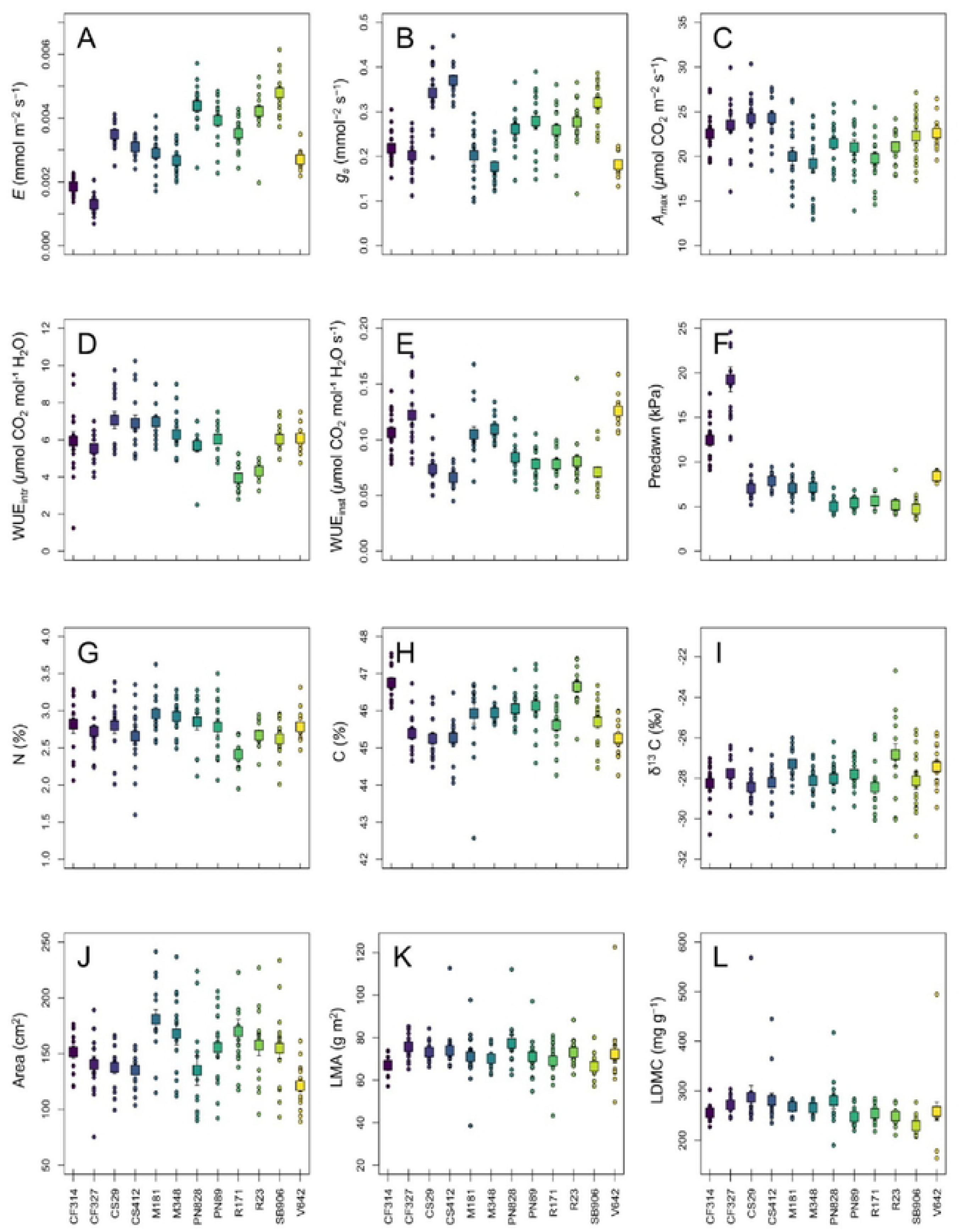
Trait variation across wine grape cultivars. Individual points (circles) represent data for individual plants, the boxes represent mean values per cultivar, and the bars represent standard errors. CF314: Cabernet Franc 314; CF327: Cabernet Franc 327; CS29: Cabernet Sauvignon 29; CS412: Cabernet Sauvignon 412; M181: Merlot 181; M348: Merlot 348; PN828: Pinot Noir 828; PN89: Pinot Noir 89; R171: Riesling 171; R23: Riesling 23; SB906: Sauvignon Blanc 906; V642: Viognier 642. A: Evapotranspiration; B: stomatal conductance; C: maximum photosynthetic capacity; D: intrinsic water use efficiency; E: instantaneous water use efficiency; F: predawn water potential; G: leaf nitrogen concentration; H: leaf carbon concentration; I: isotopic signature; J: leaf area; K: leaf mass per area; L: leaf dry matter content.

**Table 2.**
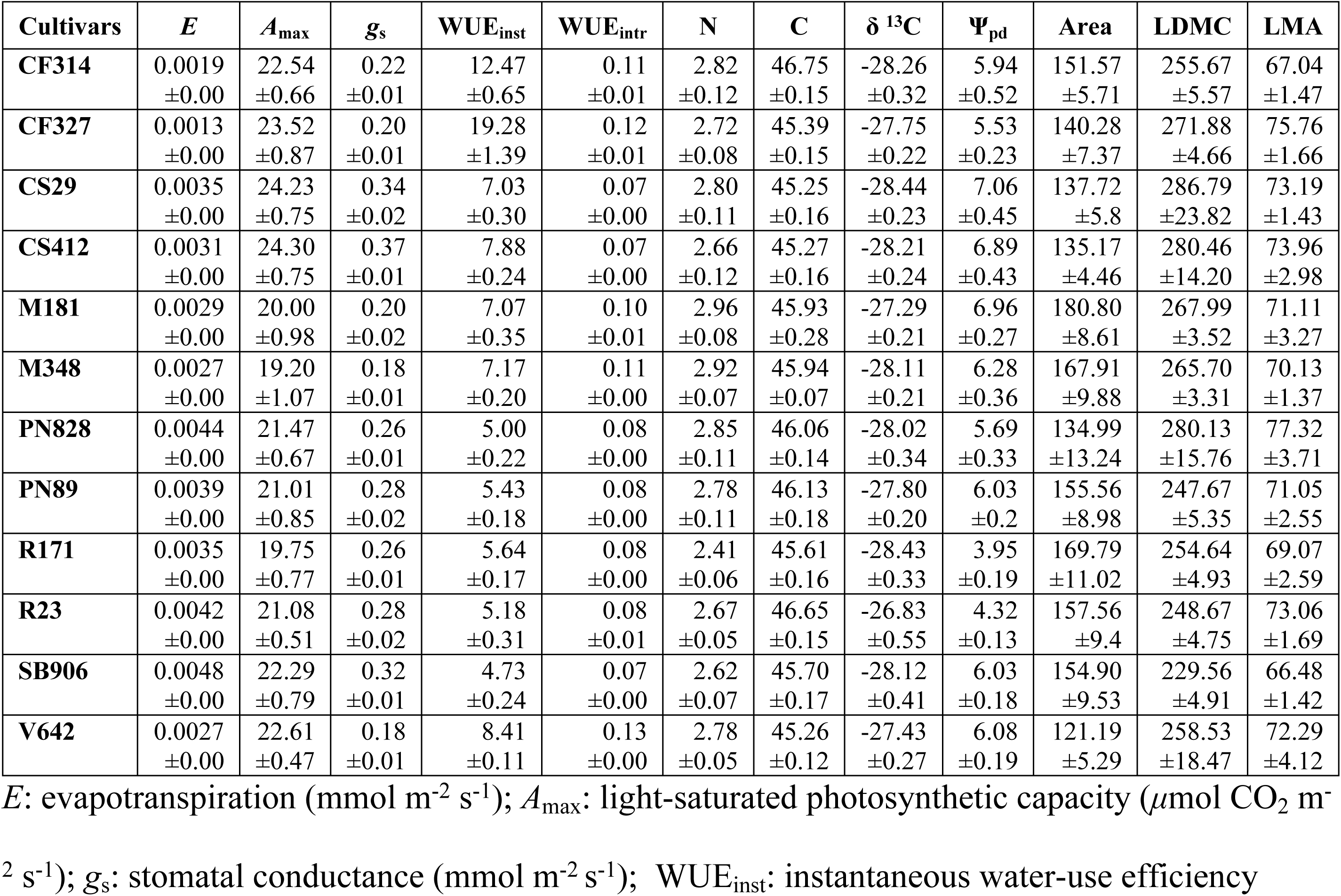
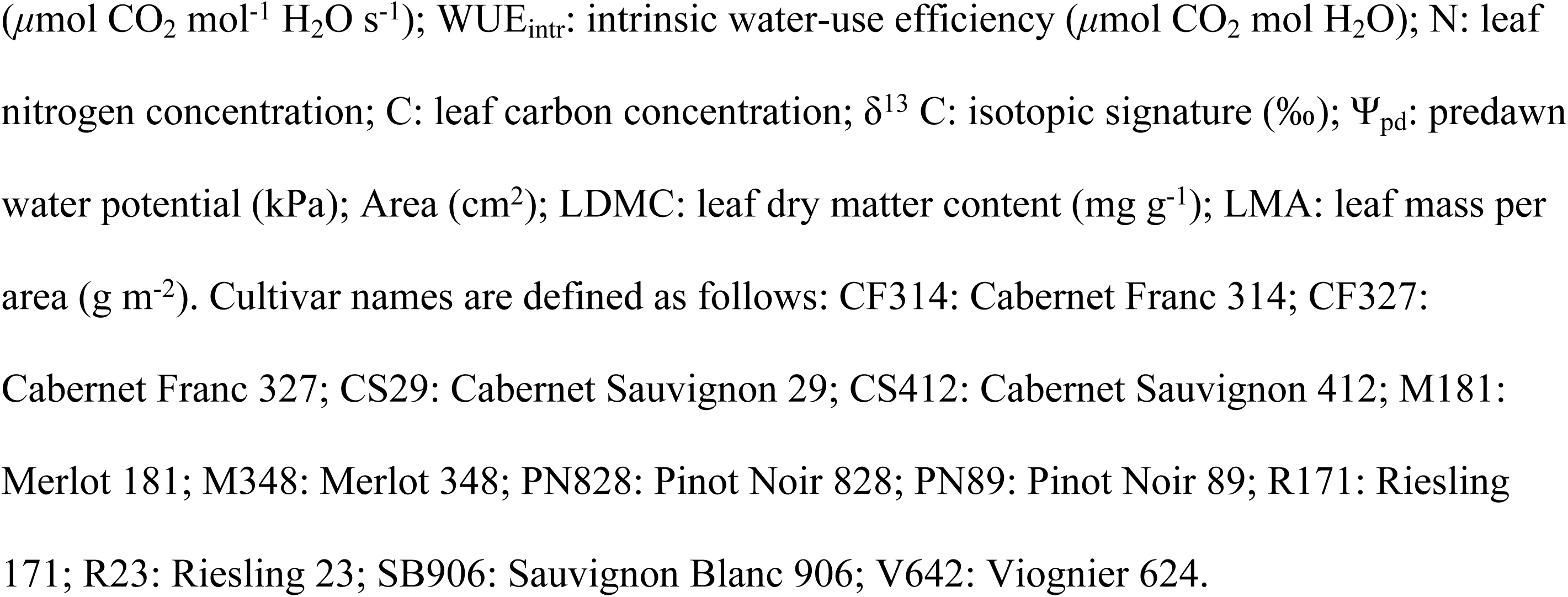
Mean trait values measured across 12 wine grape cultivars with standard error (*n*≈15 per cultivar).

| Cultivars | $E$ | $A_{\max}$ | $g_s$ | $\text{WUE}_{\text{inst}}$ | $\text{WUE}_{\text{intr}}$ | N | C | $\delta^{13}\text{C}$ | $\Psi_{\text{pd}}$ | Area | LDMC | LMA |
| --- | --- | --- | --- | --- | --- | --- | --- | --- | --- | --- | --- | --- |
| CF314 | 0.0019<br>$\pm 0.00$ | 22.54<br>$\pm 0.66$ | 0.22<br>$\pm 0.01$ | 12.47<br>$\pm 0.65$ | 0.11<br>$\pm 0.01$ | 2.82<br>$\pm 0.12$ | 46.75<br>$\pm 0.15$ | -28.26<br>$\pm 0.32$ | 5.94<br>$\pm 0.52$ | 151.57<br>$\pm 5.71$ | 255.67<br>$\pm 5.57$ | 67.04<br>$\pm 1.47$ |
| CF327 | 0.0013<br>$\pm 0.00$ | 23.52<br>$\pm 0.87$ | 0.20<br>$\pm 0.01$ | 19.28<br>$\pm 1.39$ | 0.12<br>$\pm 0.01$ | 2.72<br>$\pm 0.08$ | 45.39<br>$\pm 0.15$ | -27.75<br>$\pm 0.22$ | 5.53<br>$\pm 0.23$ | 140.28<br>$\pm 7.37$ | 271.88<br>$\pm 4.66$ | 75.76<br>$\pm 1.66$ |
| CS29 | 0.0035<br>$\pm 0.00$ | 24.23<br>$\pm 0.75$ | 0.34<br>$\pm 0.02$ | 7.03<br>$\pm 0.30$ | 0.07<br>$\pm 0.00$ | 2.80<br>$\pm 0.11$ | 45.25<br>$\pm 0.16$ | -28.44<br>$\pm 0.23$ | 7.06<br>$\pm 0.45$ | 137.72<br>$\pm 5.8$ | 286.79<br>$\pm 23.82$ | 73.19<br>$\pm 1.43$ |
| CS412 | 0.0031<br>$\pm 0.00$ | 24.30<br>$\pm 0.75$ | 0.37<br>$\pm 0.01$ | 7.88<br>$\pm 0.24$ | 0.07<br>$\pm 0.00$ | 2.66<br>$\pm 0.12$ | 45.27<br>$\pm 0.16$ | -28.21<br>$\pm 0.24$ | 6.89<br>$\pm 0.43$ | 135.17<br>$\pm 4.46$ | 280.46<br>$\pm 14.20$ | 73.96<br>$\pm 2.98$ |
| M181 | 0.0029<br>$\pm 0.00$ | 20.00<br>$\pm 0.98$ | 0.20<br>$\pm 0.02$ | 7.07<br>$\pm 0.35$ | 0.10<br>$\pm 0.01$ | 2.96<br>$\pm 0.08$ | 45.93<br>$\pm 0.28$ | -27.29<br>$\pm 0.21$ | 6.96<br>$\pm 0.27$ | 180.80<br>$\pm 8.61$ | 267.99<br>$\pm 3.52$ | 71.11<br>$\pm 3.27$ |
| M348 | 0.0027<br>$\pm 0.00$ | 19.20<br>$\pm 1.07$ | 0.18<br>$\pm 0.01$ | 7.17<br>$\pm 0.20$ | 0.11<br>$\pm 0.00$ | 2.92<br>$\pm 0.07$ | 45.94<br>$\pm 0.07$ | -28.11<br>$\pm 0.21$ | 6.28<br>$\pm 0.36$ | 167.91<br>$\pm 9.88$ | 265.70<br>$\pm 3.31$ | 70.13<br>$\pm 1.37$ |
| PN828 | 0.0044<br>$\pm 0.00$ | 21.47<br>$\pm 0.67$ | 0.26<br>$\pm 0.01$ | 5.00<br>$\pm 0.22$ | 0.08<br>$\pm 0.00$ | 2.85<br>$\pm 0.11$ | 46.06<br>$\pm 0.14$ | -28.02<br>$\pm 0.34$ | 5.69<br>$\pm 0.33$ | 134.99<br>$\pm 13.24$ | 280.13<br>$\pm 15.76$ | 77.32<br>$\pm 3.71$ |
| PN89 | 0.0039<br>$\pm 0.00$ | 21.01<br>$\pm 0.85$ | 0.28<br>$\pm 0.02$ | 5.43<br>$\pm 0.18$ | 0.08<br>$\pm 0.00$ | 2.78<br>$\pm 0.11$ | 46.13<br>$\pm 0.18$ | -27.80<br>$\pm 0.20$ | 6.03<br>$\pm 0.2$ | 155.56<br>$\pm 8.98$ | 247.67<br>$\pm 5.35$ | 71.05<br>$\pm 2.55$ |
| R171 | 0.0035<br>$\pm 0.00$ | 19.75<br>$\pm 0.77$ | 0.26<br>$\pm 0.01$ | 5.64<br>$\pm 0.17$ | 0.08<br>$\pm 0.00$ | 2.41<br>$\pm 0.06$ | 45.61<br>$\pm 0.16$ | -28.43<br>$\pm 0.33$ | 3.95<br>$\pm 0.19$ | 169.79<br>$\pm 11.02$ | 254.64<br>$\pm 4.93$ | 69.07<br>$\pm 2.59$ |
| R23 | 0.0042<br>$\pm 0.00$ | 21.08<br>$\pm 0.51$ | 0.28<br>$\pm 0.02$ | 5.18<br>$\pm 0.31$ | 0.08<br>$\pm 0.01$ | 2.67<br>$\pm 0.05$ | 46.65<br>$\pm 0.15$ | -26.83<br>$\pm 0.55$ | 4.32<br>$\pm 0.13$ | 157.56<br>$\pm 9.4$ | 248.67<br>$\pm 4.75$ | 73.06<br>$\pm 1.69$ |
| SB906 | 0.0048<br>$\pm 0.00$ | 22.29<br>$\pm 0.79$ | 0.32<br>$\pm 0.01$ | 4.73<br>$\pm 0.24$ | 0.07<br>$\pm 0.00$ | 2.62<br>$\pm 0.07$ | 45.70<br>$\pm 0.17$ | -28.12<br>$\pm 0.41$ | 6.03<br>$\pm 0.18$ | 154.90<br>$\pm 9.53$ | 229.56<br>$\pm 4.91$ | 66.48<br>$\pm 1.42$ |
| V642 | 0.0027<br>$\pm 0.00$ | 22.61<br>$\pm 0.47$ | 0.18<br>$\pm 0.01$ | 8.41<br>$\pm 0.11$ | 0.13<br>$\pm 0.00$ | 2.78<br>$\pm 0.05$ | 45.26<br>$\pm 0.12$ | -27.43<br>$\pm 0.27$ | 6.08<br>$\pm 0.19$ | 121.19<br>$\pm 5.29$ | 258.53<br>$\pm 18.47$ | 72.29<br>$\pm 4.12$ |

$E$ : evapotranspiration ( $\text{mmol m}^{-2} \text{ s}^{-1}$ ); $A_{\max}$ : light-saturated photosynthetic capacity ( $\mu\text{mol CO}_2 \text{ m}^{-2} \text{ s}^{-1}$ ); $g_s$ : stomatal conductance ( $\text{mmol m}^{-2} \text{ s}^{-1}$ ); $\text{WUE}_{\text{inst}}$ : instantaneous water-use efficiency
( $\mu\text{mol CO}_2 \text{ mol}^{-1} \text{ H}_2\text{O s}^{-1}$ ); $\text{WUE}_{\text{intr}}$ : intrinsic water-use efficiency ( $\mu\text{mol CO}_2 \text{ mol H}_2\text{O}$ ); N: leaf nitrogen concentration; C: leaf carbon concentration; $\delta^{13} \text{ C}$ : isotopic signature (‰); $\Psi_{\text{pd}}$ : predawn water potential (kPa); Area ( $\text{cm}^2$ ); LDMC: leaf dry matter content ( $\text{mg g}^{-1}$ ); LMA: leaf mass per area ( $\text{g m}^{-2}$ ). Cultivar names are defined as follows: CF314: Cabernet Franc 314; CF327: Cabernet Franc 327; CS29: Cabernet Sauvignon 29; CS412: Cabernet Sauvignon 412; M181: Merlot 181; M348: Merlot 348; PN828: Pinot Noir 828; PN89: Pinot Noir 89; R171: Riesling 171; R23: Riesling 23; SB906: Sauvignon Blanc 906; V642: Viognier 624.

We also found significant variation in traits related to plant-water relations, with WUE_inst_ varying between 4.73±0.24 and 8.41±0.11 *μ*mol CO_2_ mol^-^¹ H_2_O. There was also wide variation in the WUE_intr_ values for CF314 and CF327 (12.47±0.65 and 19.28±1.39 *μ*mol CO_2_ mol^-^¹ H_2_O, respectively), indicating an increased ability of the CF varieties to retain water while still maintaining their photosynthetic capacity. Both leaf C and N were relatively stable across the cultivars, with limited variation (Figure 1). Across cultivars, δ ^13^C also varied little, with a majority of mean values clustering around approximately −28‰, with R23 presenting the lowest average value (−26.83±0.55‰), demonstrating potentially lower long-term WUE.

Morphological traits, including leaf area, LDMC, and LMA, had wider variation across cultivars. Specifically, M181 and M348 on average had the highest leaf area, while V642 and CS412 expressed the smallest average leaf areas (121.2±5.3 and 135.2±4.5 cm^2^, respectively; Figure 2). LDMC was highest in the CS cultivars, which suggests that these leaves are denser than the other cultivars. LMA followed a similar trend; the CS cultivars had among the highest LMA values along with PN828, ranging from 73.2±1.4-77.3±3.7 g m^-2^. The differences noted here highlight the structural variations among the cultivars that reflect potential adaptation strategies.

**Figure 2.**
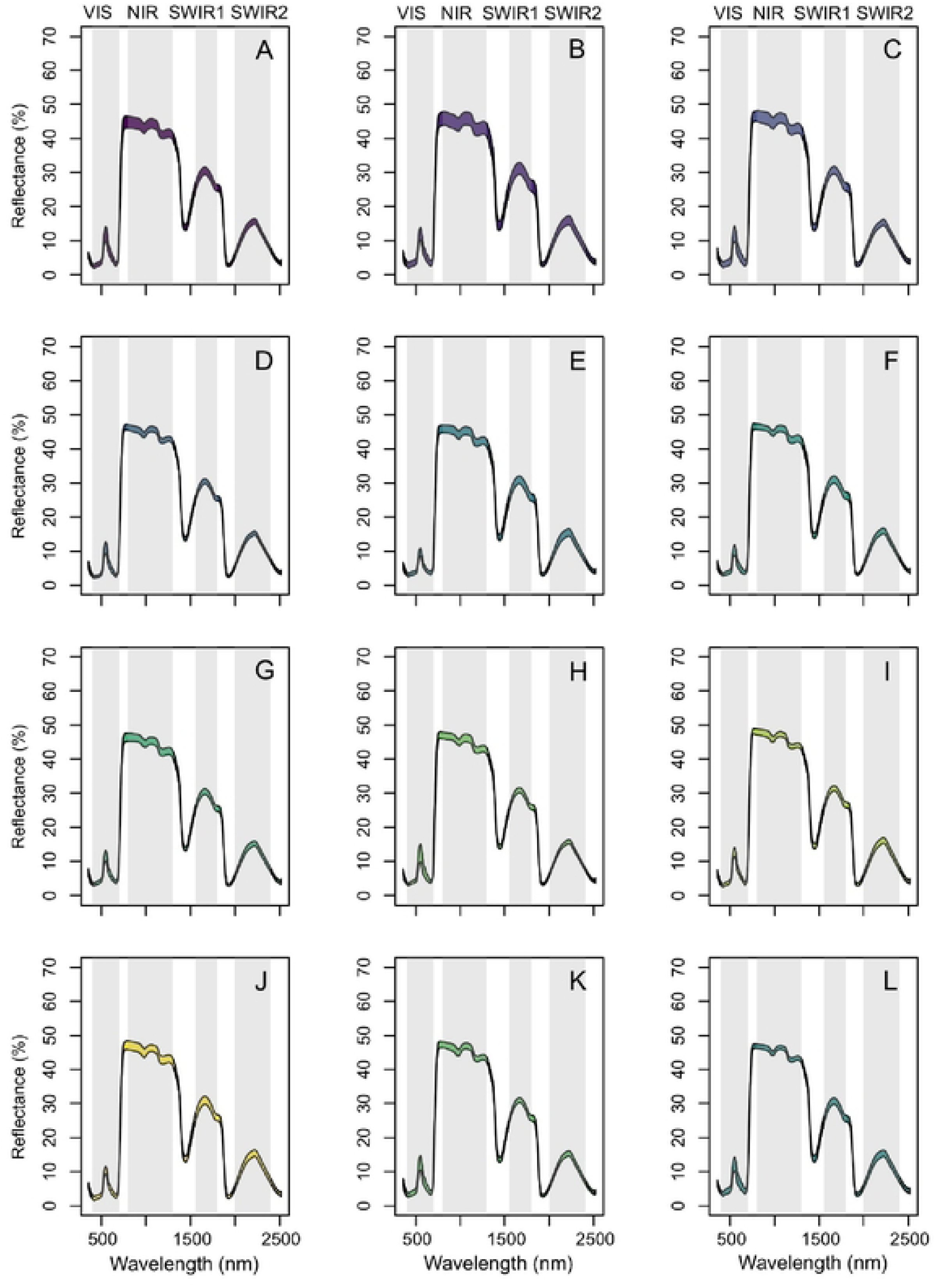
Spectra reflectance quantiles for 12 wine grape cultivars with key wavebands highlighted by the grey bars. VIS: visible light spectrum (380–780 nm); NIR: near-infrared (780–1400 nm); SWIR1: shortwave infrared radiation 1 (1570–1650 nm); SWIR2: shortwave infrared radiation 2 (2000–2290 nm). A: Cabernet Franc 314; B: Cabernet Franc 327; C: Cabernet Sauvignon 29; D: Cabernet Sauvignon 412; E: Merlot 181; F: Merlot 348; G: Pinot Noir 828; H: Pinot Noir 89; I: Riesling 171; J: Riesling 23; K: Sauvignon Blanc 906; L: Viognier 642.

### 3.3 Reflectance Spectra and PLSR Models

Figure 2 presents the spectra quantiles for all 12 wine grape cultivars (total *n*=178 leaves) with the prominent wavebands highlighted. There were no noteworthy deviations in the spectra, except for two leaves that were removed due to measurement errors. The thickness of each spectral band corresponds to the variation in the data—with thicker bands indicating higher variability in reflectance spectra—as seen in Cabernet cultivars (panels A–C: CF314, CF327, CS29). As presented in Table 3 and visualised in Figure 3, PLSR models parameterized under random data splits yielded, on average, an increase in the number of statistically significant model components, resulting in stronger predictive models. Specifically, PLSR models with the best fits in the random split were for *E* and *A*_max_, with PLSR *r*^2^ values of 0.28 and 0.33, respectively, with corresponding RMSE values of 0% and 4.84%. Additionally, LDMC had the strongest fit with an *r*^2^ value of 0.52. The models with the weakest fit include *g_s_*, WUE_inst_, and LMA, indicated by the negative *r^2^* values.

**Figure 3.**
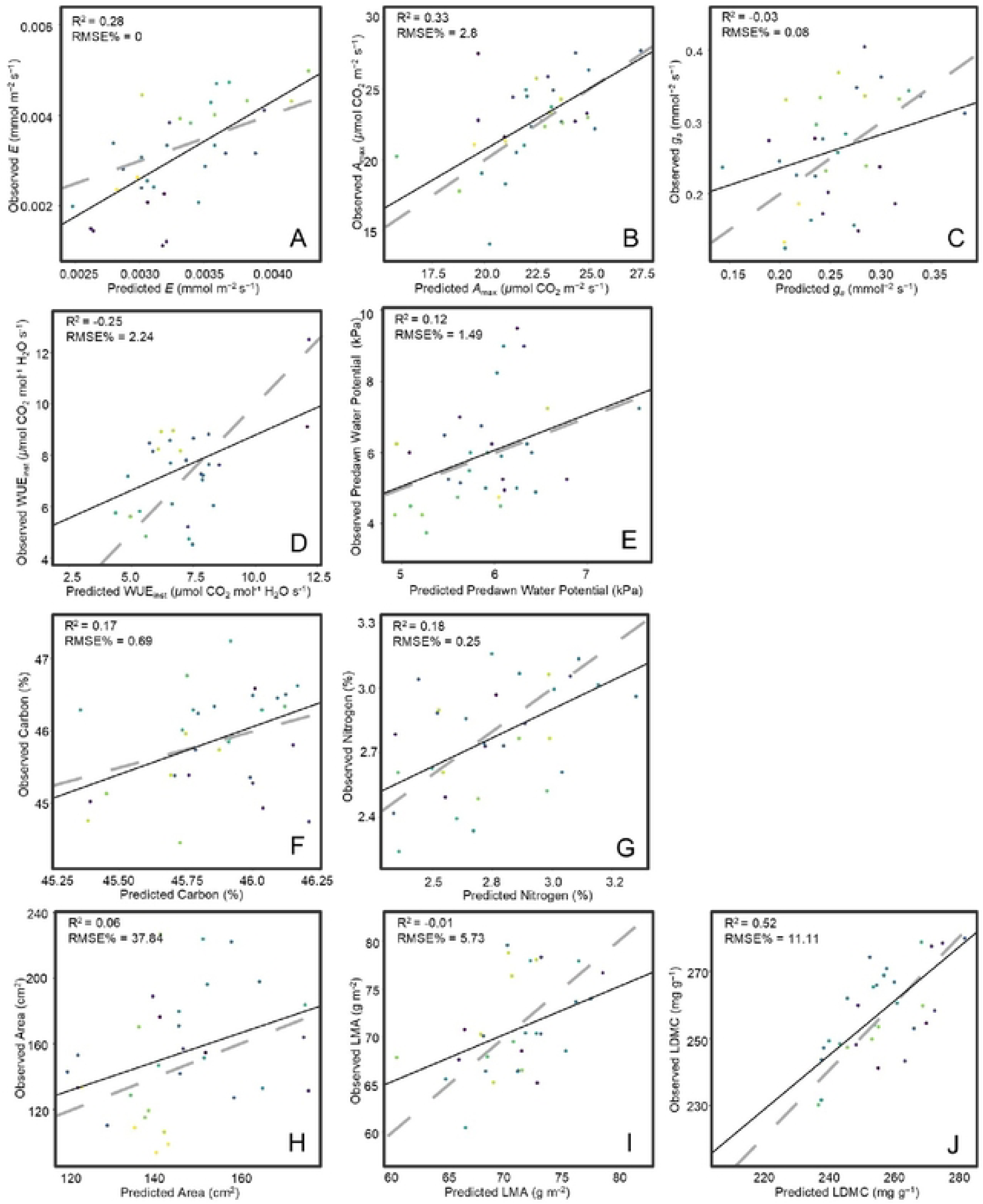
Results of partial least squares regression (PLSR) models predicting 10 leaf functional traits in 12 wine grape cultivars. Here, the data points used to validate the models fitted to a set of calibration data points are presented. Calibration and validation datasets were selected based on a “random” data split. The dashed line represents a 1:1 relationship, and the solid black line represents the linear regression model fits between the observed and predicted variables.

**Table 3.** Partial least squares regression model fits evaluating reflectance spectra to explain trait variation in leaves across 12 wine grape cultivars, with the results including a division of the data into calibration (80%) and validation (20%) datasets. The results from the “cultivar” approach include calibration/ validation data with approximately equal proportions of observations from all cultivars, whereas the “random” approach randomizes this division.

| Data Split | Trait | $n_{\text{obs}}$ | $n_{\text{cal}}$ | $n_{\text{val}}$ | $n_{\text{comp}}$ | RMSE | $r^2$ | %RMSE |
| --- | --- | --- | --- | --- | --- | --- | --- | --- |
| Cultivar | $E$ | 173 | 137 | 36 | 3 | 0.00 | 0.19 | 0.00 |
| | $g_s$ | 173 | 137 | 36 | 2 | 0.14 | -0.02 | 0.08 |
| | $WUE_{\text{inst}}$ | 165 | 130 | 35 | 4 | 0.51 | 0.12 | 0.31 |
| | $WUE_{\text{intr}}$ | 172 | 136 | 36 | 2 | 0.50 | 0.09 | 0.29 |
| | $N$ | 163 | 127 | 36 | 3 | 0.46 | 0.35 | 0.28 |
| | $C$ | 162 | 126 | 36 | 1 | 1.15 | 0.14 | 0.71 |
| | $\Psi_{\text{pd}}$ | 172 | 136 | 36 | 4 | 2.67 | -0.02 | 1.55 |
|  | Area | 168 | 132 | 36 | 1 | 50.20 | 0.22 | 29.88 |
|  | LDMC | 162 | 126 | 36 | 10 | 32.97 | 0.42 | 20.35 |
|  | LMA | 161 | 125 | 36 | 5 | 8.21 | 0.11 | 5.1 |
| Random | $E$ | 173 | 138 | 35 | 2 | 0.00 | 0.28 | 0.00 |
| | $A_{\max}$ | 173 | 138 | 35 | 9 | 4.84 | 0.33 | 2.8 |
| | $g_s$ | 173 | 138 | 35 | 8 | 0.14 | -0.03 | 0.08 |
| | $WUE_{\text{inst}}$ | 165 | 132 | 33 | 12 | 0.48 | -0.21 | 0.29 |
|  | **N** | 163 | 130 | 33 | 3 | 0.41 | 0.18 | 0.25 |
|  | **C** | 162 | 129 | 33 | 2 | 1.12 | 0.17 | 0.69 |
|  | **$\Psi_{pd}$** | 172 | 137 | 35 | 3 | 2.56 | 0.12 | 1.49 |
|  | **Area** | 168 | 134 | 34 | 10 | 63.57 | 0.06 | 37.84 |
|  | **LDMC** | 162 | 129 | 33 | 11 | 18.00 | 0.52 | 11.11 |
|  | **LMA** | 161 | 128 | 33 | 5 | 9.23 | -0.01 | 5.73 |

$n_{obs}$ : total number of observations; $n_{val}$ : total number of validation observations; $n_{comp}$ : number of components; RSME: root mean square error; RSME%: percent root mean square error (validation data); $E$ : evapotranspiration ( $\text{mmol m}^{-2} \text{s}^{-1}$ ); $A_{max}$ : light-saturated photosynthetic capacity ( $\mu\text{mol CO}_2 \text{m}^{-2} \text{s}^{-1}$ ); $g_s$ : stomatal conductance ( $\text{mmol m}^{-2} \text{s}^{-1}$ ); $\text{WUE}_{inst}$ : instantaneous water-use efficiency ( $\mu\text{mol mol m}^{-2} \text{s}^{-1}$ ); $\text{WUE}_{intr}$ : intrinsic water-use efficiency ( $\mu\text{mol mol}^{-1}$ ); N: leaf nitrogen concentration; C: leaf carbon concentration; $\Psi_{pd}$ : predawn water potential (kPa); Area ( $\text{cm}^2$ ); LDMC: leaf dry matter content ( $\text{mg g}^{-1}$ ); LMA: leaf mass per area ( $\text{g m}^{-2}$ ). Cultivar split did not successfully model $A_{max}$ , and the random split was unsuccessful for $\text{WUE}_{intr}$ .

For every measured plant trait, variable importance in projection (VIP) scores were determined to assess the importance of individual predictor variables in the PLSR output, indicating which wavebands had the greatest influence on the output. In total, all models had VIP peaks at ∼750 nm, which is near the edge of the NIR range, indicating this waveband is highly important for predicting the response variables. In addition, LMA, LDMC, and area had VIP peaks within the visible light spectrum (500 nm), while WUE_inst_ and *g*_s_ had VIP peaks in the SWIR range between 1000-1800 nm (Figure 4).

**Figure 4.**
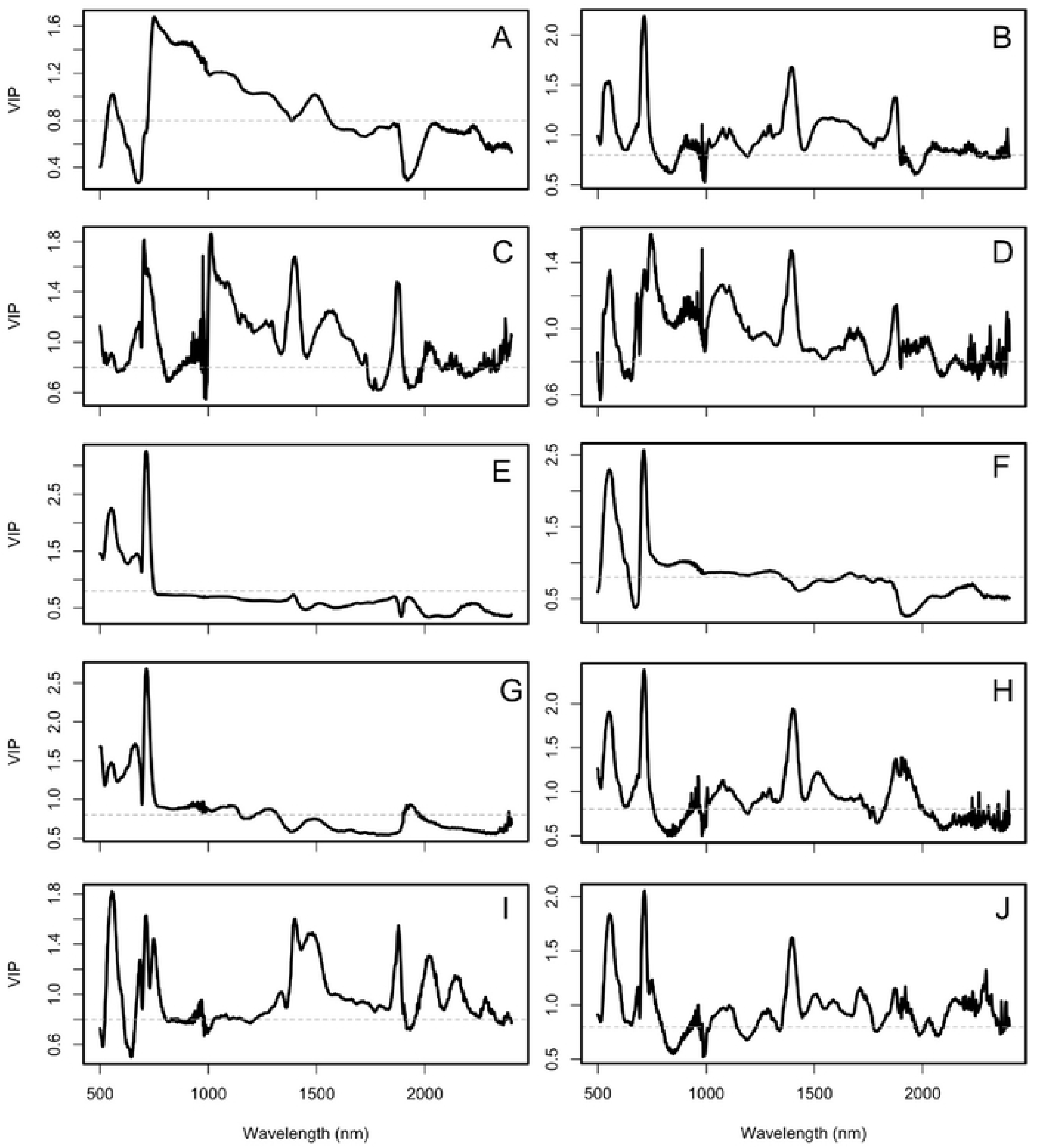
Variable influences on projection (VIP) scores associated with partial least squares regression models for the calibration datasets using a random split. The dashed line represents where the VIP score is 0.8. A: Evapotranspiration; B: maximum assimilation rate; C: stomatal conductance; D: instantaneous water-use efficiency; E: leaf nitrogen concentration; F: leaf carbon concentration; G: predawn water potential; H: area; I: leaf mass per area; J: leaf dry matter content.

## Discussion

This study contributes to the growing body of research demonstrating the application of reflectance spectroscopy and PLSR modelling for quantifying intraspecific or single-species trait variation. Previous work has established this approach to trait estimation across different species (20,21,28), with more recent studies exploring its application in wine grapes (26). More recently, the importance of understanding and quantifying intraspecific trait variation as a driver for plant- and ecosystem-level resilience (16,31) has gained increased attention, motivating an interest in using reflectance spectroscopy and PLSR modelling for functional trait characterisation, especially in agroecosystems and crop breeding programs (20). Here, we explored the extent and sources of functional trait variation across a diverse genetic gradient of wine grape cultivars using both field-based research methods and leaf-level spectral data.

Our findings demonstrate that plant traits varied across cultivars, indicating a strong influence of winegrape genotypes on phenotypic trait expression. This variation was most evident in the resource acquisition and water-use traits, with less distinctive variation in the leaf chemical and morphological traits. Similar patterns have been documented in other single-species studies, and our results align with these findings, confirming that there are both environmental and genetic bases for trait variation, and measuring such variation is critical for predicting plant responses to change. In line with other studies (26,32) we also detected that the ∼750 nm range had the strongest influence on trait estimation for a majority of the measured traits, with the lower SWIR range wavebands having an impact on the physiological and water-use traits.

In our analysis, we found that PLSR models predict three distinctive physiological and morphological leaf traits with a predictive power range of 28% to 52%, for *E*, *A*_max_, and LDMC, and we had moderate success predicting four additional leaf chemical and morphological traits, including leaf C and N concentrations, Ψ_pd_, and area. Compared to other studies focusing on single-species, our PLSR models were less effective in trait prediction. For example, in similar studies, the predictive power of the PLSR models for the leaf photosynthetic traits for six winegrape varieties was much higher, ranging from 18-62% (23). The ability of the PLSR models to predict variation in leaf traits across wine grape cultivars was dependent on data splitting. By fully randomising the data split when building the calibration and validation datasets, it resulted in higher *r*^2^ and %RMSE values across 10 of the measured traits that were successfully modelled. This is likely due to the general variability and size of our dataset. When working with smaller datasets, having a non-random data split may distort or overfit the model, providing unreliable performance metrics. Thus, ahead of model calibration, data splits need to be assessed based on the general variability of the training data and how this variation is structured (21,28). Additionally, the data splitting strategy can also yield different outcomes depending on the trait assessed. Traits with lower environmental specificity may be more accurately predicted with the random split, whereas traits with more pronounced genetic specificity may be more accurately modelled using a cultivar split, as presented in Table 3.

## Conclusions

This study demonstrates that functional traits in wine grape leaves vary substantially across cultivars. Traits related to water-use and photosynthetic capacity are particularly variable, highlighting their potential use for selecting resilient cultivars for breeding programs and climate adaptation strategies. Reflectance spectroscopy and the subsequent PLSR models used show significant promise for detecting and estimating specific plant traits; however, these models will benefit from enhanced calibration and using larger datasets to increase accuracy and performance. With further calibration, these models can be scaled up to assess the successful plant traits using reflectance measurements captured through hyperspectral imaging, to link in-field data with landscape-level crop spectral data (33).

## Acknowledgements

The authors would also like to thank the Natural Sciences and Engineering Research Council of Canada, University of Toronto Scarborough. The authors acknowledge Emily Young and Kale Vicario for their assistance with field data collection, and Camila Villa-Escobar and Rachael Harman-Denhoed for their assistance with the sample preparation and processing.

## References

1. Rezaei E, Webber H, Asseng S, Boote K, Durand JL, Ewert F, et al. Climate change impacts on crop yields. Nat Rev Earth Environ. 2023 Nov 14;4.

2. van Leeuwen C, Sgubin G, Bois B, Ollat N, Swingedouw D, Zito S, et al. Climate change impacts and adaptations of wine production. Nat Rev Earth Environ. 2024 Apr;5(4):258–75.

3. Seleiman MF, Al-Suhaibani N, Ali N, Akmal M, Alotaibi M, Refay Y, et al. Drought Stress Impacts on Plants and Different Approaches to Alleviate Its Adverse Effects. Plants. 2021 Feb;10(2):259.

4. Jones GV, Reid R, Vilks A. Climate, Grapes, and Wine: Structure and Suitability in a Variable and Changing Climate. In: Dougherty PH, editor. The Geography of Wine: Regions, Terroir and Techniques [Internet]. Dordrecht: Springer Netherlands; 2012 [cited 2025 Jul 31]. p. 109–33. Available from: 10.1007/978-94-007-0464-0_7

5. Annual Reports – Wine Growers Ontario [Internet]. [cited 2025 Aug 14]. Available from: https://www.winegrowersontario.ca/annual-reports/

6. Keller M. Climate Change Impacts on Vineyards in Warm and Dry Areas: Challenges and Opportunities: From the ASEV Climate Change Symposium Part 1 – Viticulture. Am J Enol Vitic [Internet]. 2023 Jul 1 [cited 2025 Jul 31];74(2). Available from: https://www.ajevonline.org/content/74/2/0740033

7. Morales-Castilla I, García de Cortázar-Atauri I, Cook BI, Lacombe T, Parker A, van Leeuwen C, et al. Diversity buffers winegrowing regions from climate change losses. Proc Natl Acad Sci. 2020 Feb 11;117(6):2864–9.

8. Bush E, Lemmen DS. Canada’s changing climate report [Internet]. Government of Canada; 2019 [cited 2025 Aug 18]. Available from: https://ostrnrcan-dostrncan.canada.ca/handle/1845/143725

9. Baltazar M, Castro I, Gonçalves B. Adaptation to Climate Change in Viticulture: The Role of Varietal Selection—A Review. Plants. 2025 Jan;14(1):104.

10. Demmings EM, Williams BR, Lee CR, Barba P, Yang S, Hwang CF, et al. Quantitative Trait Locus Analysis of Leaf Morphology Indicates Conserved Shape Loci in Grapevine. Front Plant Sci [Internet]. 2019 Nov 15 [cited 2025 Apr 7];10. Available from: https://www.frontiersin.org/journals/plant-science/articles/10.3389/fpls.2019.01373/full

11. Guo A, Zuo X, Zhang S, Hu Y, Yue P, Lv P, et al. Contrasting effects of plant inter- and intraspecific variation on community trait responses to nitrogen addition and drought in typical and meadow steppes. BMC Plant Biol. 2022 Mar 1;22(1):90.

12. Miedema Brown L, Anand M. Plant functional traits as measures of ecosystem service provision. Ecosphere. 2022 Feb;13(2):e3930.

13. Violle C, Enquist BJ, McGill BJ, Jiang L, Albert CH, Hulshof C, et al. The return of the variance: intraspecific variability in community ecology. Trends Ecol Evol. 2012 Apr 1;27(4):244–52.

14. Martin AR, Li G, Cui B, Mariani RO, Vicario K, Cathline KA, et al. A high-throughput approach for quantifying turgor loss point in grapevine. Plant Methods. 2024 Nov 24;20(1):180.

15. Nimmo V, Violle C, Entz M, Rolhauser AG, Isaac ME. Changes in crop trait plasticity with domestication history: Management practices matter. Ecol Evol [Internet]. 2023 [cited 2025 Sep 30];13(11). Available from: https://onlinelibrary.wiley.com/doi/10.1002/ece3.10690

16. Westerband AC, Funk JL, Barton KE. Intraspecific trait variation in plants: a renewed focus on its role in ecological processes. Ann Bot. 2021 Apr 1;127(4):397–410.

17. Isaac ME, Martin AR. Accumulating crop functional trait data with citizen science. Sci Rep. 2019 Oct 31;9(1):15715.

18. Pabuayon ILB, Sun Y, Guo W, Ritchie GL. High-throughput phenotyping in cotton: a review. J Cotton Res. 2019 Oct 29;2(1):18.

19. Ely KS, Burnett AC, Lieberman-Cribbin W, Serbin SP, Rogers A. Spectroscopy can predict key leaf traits associated with source-sink balance and carbon-nitrogen status. J Exp Bot. 2019 Mar 27;70(6):1789–99.

20. Kim M, Lee C, Hong S, Kim SL, Baek JH, Kim KH. High-Throughput Phenotyping Methods for Breeding Drought-Tolerant Crops. Int J Mol Sci. 2021 Jul 31;22(15):8266.

21. Kothari S, Beauchamp-Rioux R, Laliberté E, Cavender-Bares J. Reflectance spectroscopy allows rapid, accurate and non-destructive estimates of functional traits from pressed leaves. Methods Ecol Evol. 2023;14(2):385–401.

22. Cotrozzi L, Peron R, Tuinstra MR, Mickelbart MV, Couture JJ. Spectral Phenotyping of Physiological and Anatomical Leaf Traits Related with Maize Water Status1[OPEN]. Plant Physiol. 2020 Nov;184(3):1363–77.

23. Charrier G. Extrapolating physiological response to drought through step-by-step analysis of water potential. Plantae. 2020 Oct;184(2).

24. Ma WT, Yu YZ, Wang X, Gong XY. Estimation of intrinsic water-use efficiency from δ13C signature of C3 leaves: Assumptions and uncertainty. Front Plant Sci. 2023 Jan 12;13:1037972.

25. Banks JM, Hirons AD. Alternative methods of estimating the water potential at turgor loss point in Acer genotypes. Plant Methods. 2019 Apr 4;15(1):34.

26. Cui B, Mariani R, Cathline KA, Robertson G, Martin AR. Reflectance spectroscopy predicts leaf functional traits across wine grape cultivars. PLANTS PEOPLE PLANET [Internet]. 2025 [cited 2025 Jul 6];n/a(n/a). Available from: https://onlinelibrary.wiley.com/doi/abs/10.1002/ppp3.70024

27. JP (S, to 2007) DB (up, to 2002) SD (up, to 2005) DS (up, authors (src/rs.f) E, sigma) SH (Author fixed, et al. nlme: Linear and Nonlinear Mixed Effects Models [Internet]. 2025 [cited 2025 Oct 20]. Available from: https://cran.r-project.org/web/packages/nlme/index.html

28. Burnett AC, Anderson J, Davidson KJ, Ely KS, Lamour J, Li Q, et al. A best-practice guide to predicting plant traits from leaf-level hyperspectral data using partial least squares regression. J Exp Bot. 2021 Sep 30;72(18):6175–89.

29. Meireles JE, Schweiger AK, Cavender-Bares J. spectrolab: Class and Methods for Spectral Data [Internet]. 2025 [cited 2025 Oct 20]. Available from: https://cran.r-project.org/web/packages/spectrolab/index.html

30. Liland KH, Mevik BH, Wehrens R, Hiemstra P. pls: Partial Least Squares and Principal Component Regression [Internet]. 2024 [cited 2025 Oct 20]. Available from: https://cran.r-project.org/web/packages/pls/index.html

31. Martin AR, Isaac ME. Functional traits in agroecology: Advancing description and prediction in agroecosystems. J Appl Ecol. 2018;55(1):5–11.

32. Serbin SP, Singh A, Desai AR, Dubois SG, Jablonski AD, Kingdon CC, et al. Remotely estimating photosynthetic capacity, and its response to temperature, in vegetation canopies using imaging spectroscopy. Remote Sens Environ. 2015 Sep 15;167:78–87.

33. Ji F, Li F, Hao D, Shiklomanov AN, Yang X, Townsend PA, et al. Unveiling the transferability of PLSR models for leaf trait estimation: lessons from a comprehensive analysis with a novel global dataset. New Phytol. 2024;243(1):111–31.

